# Growth stage-dependent changes of the levels of keratin 1 and keratin 10 as well as skin’s green autofluorescence of the back and the ears of C57BL/6 mice under basal conditions and after UVC irradiation

**DOI:** 10.1101/2020.12.20.423664

**Authors:** Zhaoxia Yang, Mingchao Zhang, Weihai Ying

**Author notes:** Corresponding author: Weihai Ying, Ph.D., Professor, School of Biomedical Engineering and Med-X Research Institute, Shanghai Jiao Tong University, 1954 Huashan Road, Shanghai, 200030, P.R. China.

## Abstract

Our previous studies have indicated that oxidative stress and inflammation can dose-dependently induce increased skin’s green autofluorescence (AF) of mice, which results at least partially from keratin 1 (K1) cleavage. Increased green AF was also found in patients’ skin of several major diseases, which may become a novel biomarker for non-invasive diagnosis. We also found age-dependent changes of the skin’s green AF of natural populations. In this study we tested our hypothesis that there are growth stage-dependent changes of K1 and keratin 10 (K10) levels in the skin of mice, which may underlie the age-dependent changes of the skin’s green AF. We found that in the skin of both mice’s back and ears, there were growth stage-dependent changes of the levels of K1 and K10 as well as the basal green AF. The K1 and K10 levels in the back’s skin were significantly different from those in the ear’s skin. There were also growth stage-dependent changes of the UVC-induced changes of K1 and K10 levels of both the ears and the back. Collectively, our study has provided first evidence showing growth stage-dependent and differential changes of the levels of K1 and K10 as well as skin’s green AF in the back and the ears of mice under basal conditions and after UVC irradiation. These findings are valuable for understanding the age-dependent changes of the skin’s green AF of natural populations, which are also important for establishing the keratins’ AF-based method for non-invasive diagnosis of diseases.

## Introduction

Keratins play multiple significant roles in epithelium, including intermediate filament formation ^[1]^, inflammatory responses ^[2, 3]^ and cellular signaling ^[4, 5]^. The proteins are also widely used as diagnostic tumor markers ^[4]^. Keratin 1 (K1) and its heterodimer partner keratin 10 (K10) are the major keratins in the suprabasal keratinocytes of epidermis ^[6–8]^, which is a hallmark for keratinocyte differentiation ^[9]^. Increasing evidence has suggested new biological functions of keratins, e.g., K1 is also an integral component of the multiprotein kininogen receptor of endothelial cells ^[10, 11]^. There are several K1 mutations- and K10 mutations-associated diseases ^[12–15]^. However, so far there has been significant deficiency in the information regarding the distribution of K1 and K10 as well as growth stage-dependent changes of K1 and K10 in the body, which is required for elucidating the pathological changes of the K1 mutation- and K10 mutation-associated diseases.

Human autofluorescence (AF) has shown great promise for non-invasive diagnosis of diabetes ^[16]^ and cancer ^[17]^. Keratins, together with melanin, NADH and FAD, are major epidermal fluorophores ^[18]^. Our previous studies have indicated that oxidative stress and inflammation can dose-dependently induce increased skin’s green autofluorescence (AF) of mice, which results at least partially from K1 cleavage ^[19]^. In multiple locations of the skin, the patients of each of the diseases we have studied, including AIS ^[20]^, myocardial infarction ^[21]^, stable coronary artery disease ^[21]^, Parkinson’s disease ^[22]^ and lung cancer ^[23]^, had significant increases in skin’s green AF, compared with age-matched healthy controls.

Our previous study found age-dependent changes of the skin’s green AF of natural populations ^[24]^, while the mechanisms underlying these changes have remained unknown. In this study we tested our hypothesis that there are growth stage-dependent changes of K1 and K10 levels in the skin of mice, which may underlie the age-dependent changes of the skin’s green AF. Our study has provided first evidence showing growth stage-dependent and differential changes of the levels of K1 and K10 as well as skin’s green AF in the back and the ears of mice under basal conditions and after UVC irradiation.

## Materials and Methods

### Materials

All chemicals were purchased from Sigma (St. Louis, MO, USA) except where noted.

### Animal Studies

Male C57BL/6Slac mice SPF grade were purchased from JSJ Laboratory (Shanghai, China). All of the animal protocols were approved by the Animal Study Committee of the School of Biomedical Engineering, Shanghai Jiao Tong University.

### Exposures of UV radiation

As described previously, an UVC lamp (TUV 25W/G25 T8, Philips, Hamburg, Germany) was used as the UV sources in our experiments. C57BL/6 mice were used for UVC treatment. After the mice were briefly anesthetized with 3.5% (w/v) chloral hydrate (1 ml /100 g), the backs and ears of the mice were exposed to UV lamps. The power density of UVC was 0.55 ± 0.05mW/cm^2^, measured by a UVC detector (ST-512, UVC, SENTRY OPTRONICS CORP., Taiwan, China).

### Imaging of mice’s epidermal AF

The epidermal AF of the backs and ears of the mice was imaged by a two-photon fluorescence microscope (A1 plus, Nikon Instech Co., Ltd., Tokyo, Japan), with the excitation wavelength of 488 nm and the emission wavelength of 500 - 530 nm. The AF was quantified by the following approach: Sixteen spots with the size of approximately 100 × 100 μm^2^ on the scanned images were selected randomly. After the average AF intensities of each layer were calculated, the sum of the average AF intensities of all layers of each spot was calculated, which is defined as the AF intensity of each spot. If the value of average AF intensity of certain layer is below 45, the AF signal of the layer is deemed background noise, which is not counted into the sum.

### Western blot assays

The lysates of the skin were centrifuged at 12,000 g for 20 min at 4 °C. The protein concentrations of the samples were quantified using BCA Protein Assay Kit (Pierce Biotechonology, Rockford, IL, USA). As described previously (17), a total of 50 μg of total protein was electrophoresed through a 10% SDS-polyacrylamide gel, which were then electrotransferred to 0.45 μm nitrocellulose membranes (Millipore, CA, USA). The blots were incubated with a monoclonal Anti-Cytokeratin 1 (ab185628, Abcam, Cambridge, UK) (1:4000 dilution) or actin (1:1000, sc-58673, Santa Cruz Biotechnology, Inc., Dallas, TX, USA) with 0.05% BSA overnight at 4°C, then incubated with HRP conjugated Goat Anti-Rabbit IgG (H+L) (1:4000, Jackson ImmunoResearch, PA, USA) or HRP conjugated Goat Anti-mouse IgG (1:2000, HA1006, HuaBio, Zhejiang Province, China). An ECL detection system (Thermo Scientific, Pierce, IL, USA) was used to detect the protein signals. The intensities of the bands were quantified by densitometry using Image J.

### Immunofluorescence assays

In some studies where noted, after the backs and ears of C57BL/6 mouse were irradiated with UVC, the backs and ears were collected, which were imaged under a confocal fluorescence microscope (TCS SP5Ⅱ, Lecia, Wetzlar, Germany). The excitation wavelength was 488 nm and the emission wavelength was 500-550 nm. The AF was quantified by using the following protocol: Sixteen spots with the size of approximately 100 × 100 μm^2^ on the scanned images were selected randomly. The average AF intensities of the 16 spots were quantified. The average value of the 16 AF intensities was defined as the AF intensity of the sample.

### Statistical analyses

All data are presented as mean ± SEM. Data were assessed by one-way ANOVA, followed by Student - Newman - Keuls post hoc test. P values less than 0.05 were considered statistically significant.

## Results

### 1. There are growth stage-dependent changes of the levels of K1 and K10 as well as skin’s green AF of the back and the ears of C57BL/6 mice under basal conditions

We determined the levels of K1 and K10 in both the back’s skin and the ears’ skin of C57BL/6 mouse, showing obvious growth stage-dependent changes of the proteins in the skin of both the back and the ears. Our Western blot assays showed that the K1 levels of the back’s skin were significantly increased from the age of 1 d to 1 w, which were then decreased at both the age of 3 w and 10 w (Figs. 1A and 1B). In contrast, the K10 levels of the back’s skin were continuously increased from the age of 1 d to the age of 1 w, 3 w and 10 w (Figs. 1A and 1C). Our Western blot assays also showed that the K1 level of the ear’s skin at the age of 10 w was significantly lower than that of the ears at the age of 3 w (Figs. 1D and 1E). In contrast, the K10 level of the ear’s skin at the age of 10 w was significantly higher than that of the ear’s skin at the age of 3 w (Figs. 1D and 1F).

**Fig.1.**
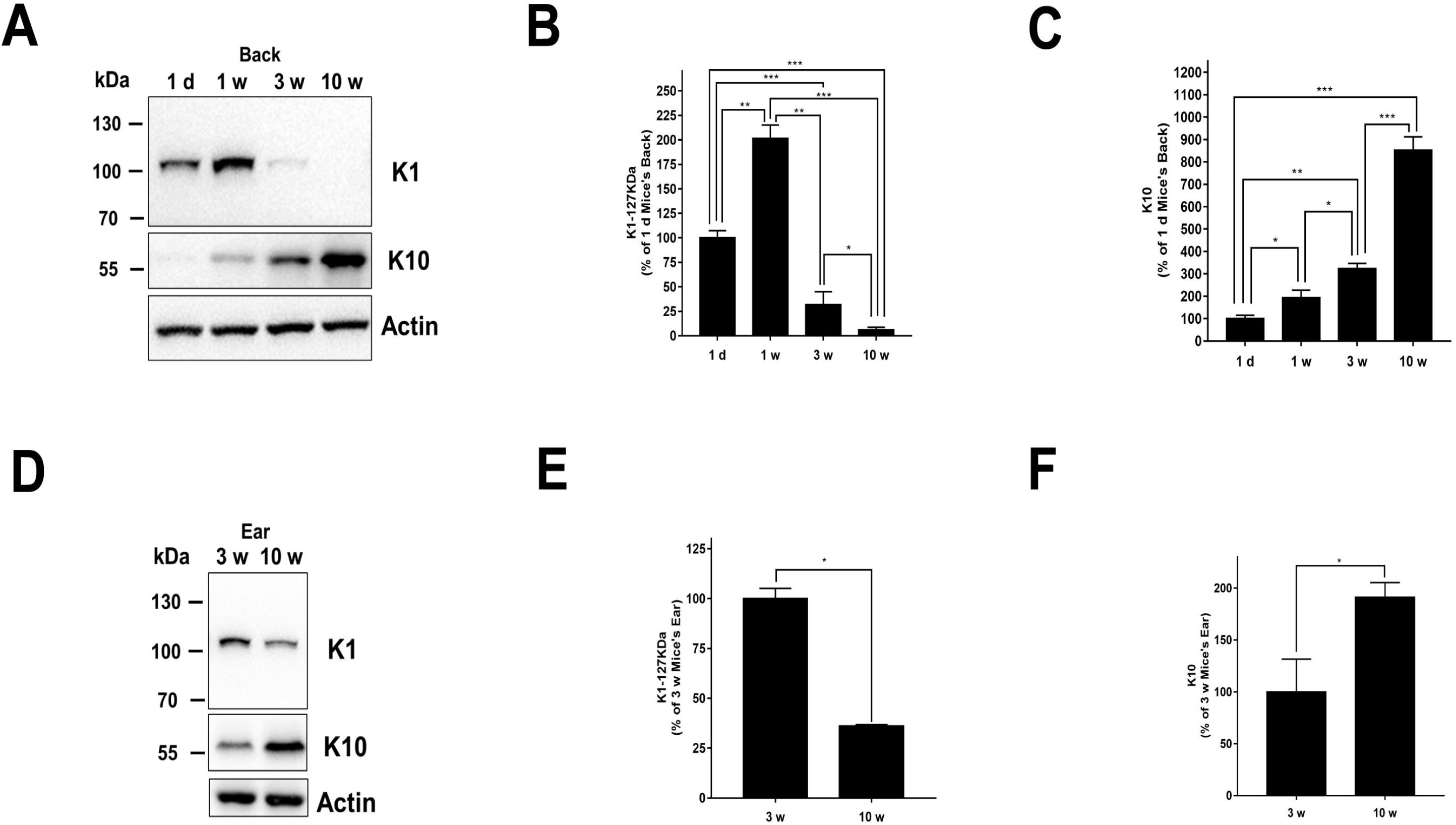
There were growth stage-dependent changes of the levels of K1 and K10 of the back and the ears of C57BL/6 mice under basal conditions. (A) Western blot assays showed that the K1 and K10 levels of the back’s skin of the mice at the age of 1 d, 1 w, 3 w, 10 w were significantly different from each other. (B) Quantifications of the Western blot showed that the K1 levels of the back’s skin were increased from the age of 1 d to 1 w, which were then decreased at the age of 3 w and 10 w. N = 9; *; *p* < 0.05; **; *p* < 0.01; ***; *p* < 0.001. (C) Quantifications of the Western blot showed that the K10 levels of the back’s skin were continuously increased from the age of 1 d to the age of 1 w, 3 w and 10 w. N = 9; *; *p* < 0.05; **; *p* < 0.01; ***; *p* < 0.001. (D) Our Western blot assays showed that the K1 and K10 levels of the ear’s skin changed from the age of 3 w to the age of 10 w. (E) Quantifications of the Western blot showed that the K1 levels of the ear’s skin at the age of 10 w was significantly lower than that of the ears at the age of 3 w. N = 9; *; *p* < 0.05. (F) Quantifications of the Western blot showed that the K10 level of the ear’s skin at the age of 10 w was significantly higher than that of the ear’s skin at the age of 3 w. N = 9; *; *p* < 0.05.

Using anti-K1 antibody, our immunostaining assays on the back’s skin of the mice at the age of 1d, 1 w, 3 w, and 10 w showed that the K1 levels changed in the similar pattern as those observed in our Western blot assays (Figs. 2A). Our immunostaining assays on the ear’s skin of the mice at the age of 3 w and 10 w also showed that the K1 levels changed in the similar pattern as those observed in our Western blot assays (Figs. 2B).

**Fig.2.**
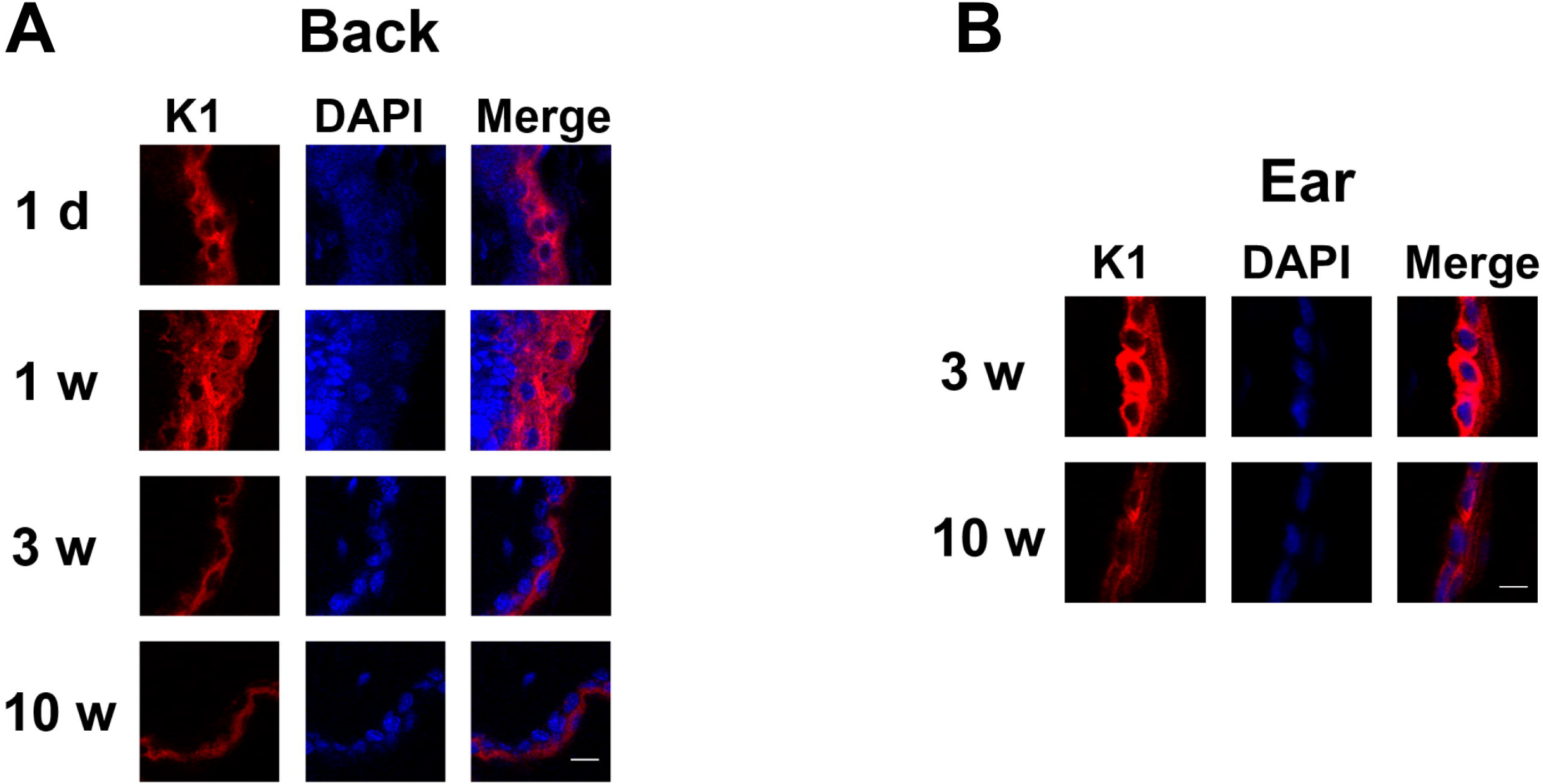
Immunostaining assays using anti-K1 antibody on the back’s skin and ear’s skin of the mice at the various growth stages showed that both K1 and K10 levels changed in the similar patterns as those observed in our Western blot assays. (A) Immunostaining assays using anti-K1 antibody on the back’s skin of the mice at the age of 1 d, 1 w, 3 w, 10 w showed that the K1 levels changed in the similar patterns as those observed in our Western blot assays. N = 3 - 6. Scale bar = 5 μm. (B) Immunostaining assays using anti-K1 antibody on the ear’s skin of the mice at the age of 3 w and 10 w showed that the K1 levels changed in the similar patterns as those observed in our Western blot assays. N = 3 - 6. Scale bar = 5 μm.

For the mice at the age of 3 w and 10 w, the K1 levels of the ears’ skin of the mice were significantly higher than those of the back’ skin (Figs. 3A and 3B). In contrast, for the mice at the age of 3 w and 10 w, the K10 levels of the ear’s skin were significantly lower than those of the back’s skin (Figs. 3A and 3C).

**Fig.3.**
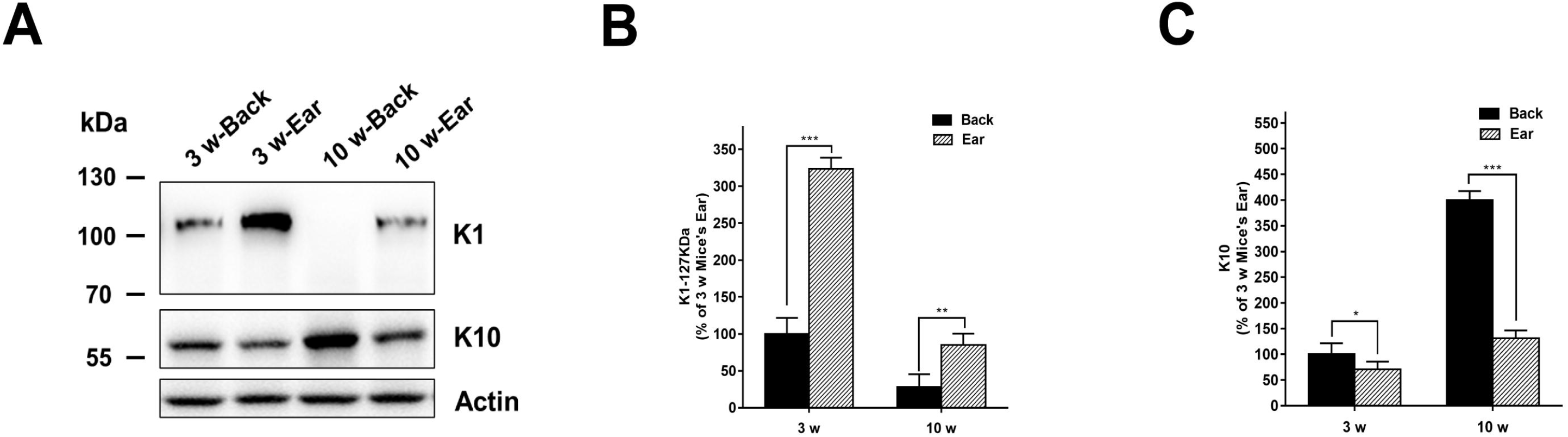
The K1 and K10 levels of the back’s skin were different from those of the skin’s skin of the mice. (A) Western blot assays showed that the K1 and K10 levels of the back’s skin were different from those of the skin’s skin of the mice. (B) For the mice at the age of 3 w and 10 w, the K1 levels of the ears’ skin of the mice were significantly higher than those of the back’ skin. N = 9; **; *p* < 0.01; ***; *p* < 0.001. (C) For the mice at the age of 3 w and 10 w, the K10 levels of the ear’s skin were significantly lower than those of the back’s skin. N = 9; *; *p* < 0.05; ***; *p* < 0.001.

We determined the basal green AF of the back’s skin and the ear’s skin of the mice. The basal AF intensity of the back’s skin at the age of 1 d was significantly higher than that at the age of 1 w, 3 w and 10 w (Figs. 4A and 4B). In contrast, the basal AF intensity of the ear’s skin at the age of 10 w was significantly higher than that at the age of 3 w (Figs. 4A and 4C).

**Fig.4.**
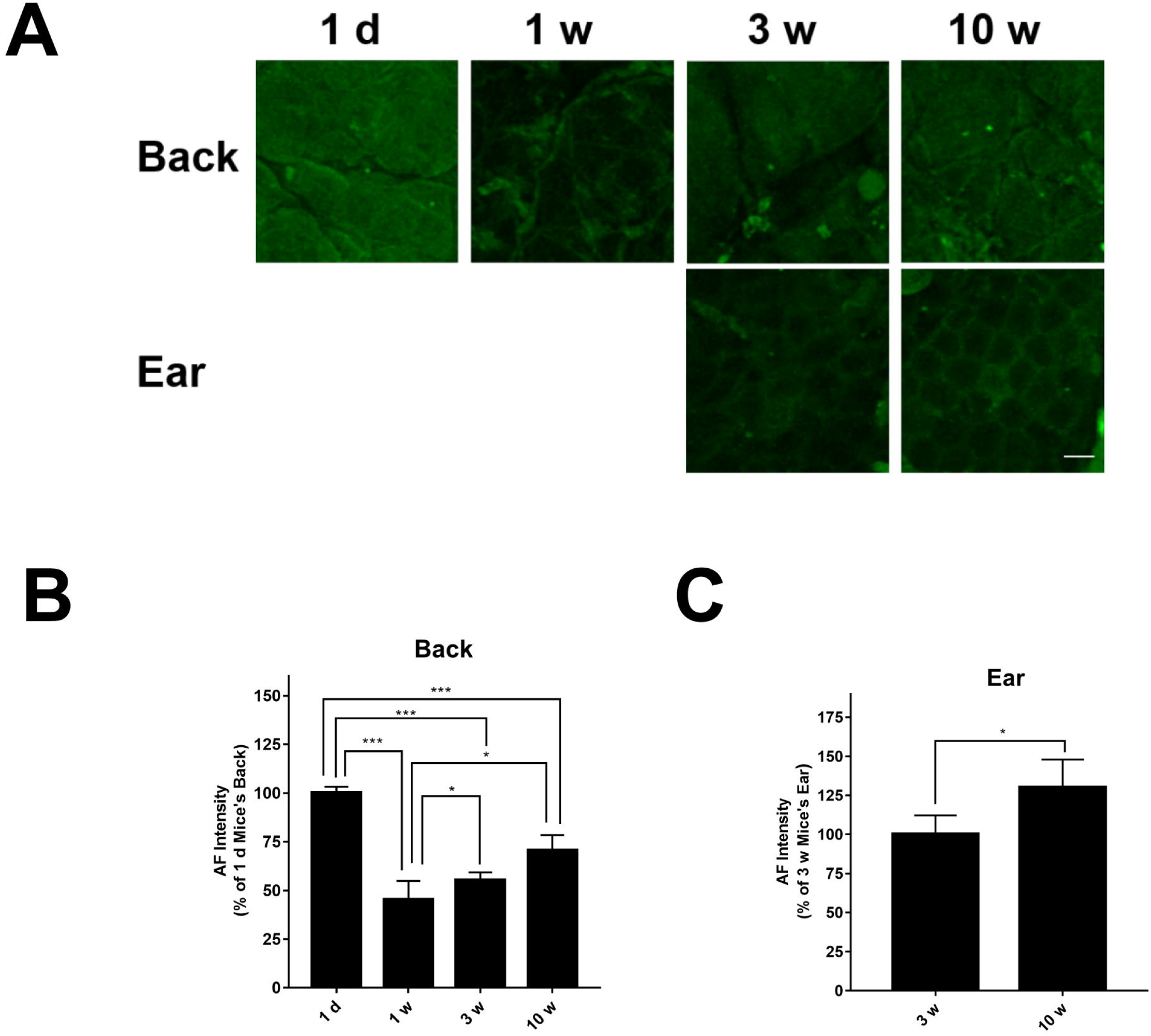
There were growth stage-dependent changes of the basal green AF of the back’s skin and the ear’s skin of the mice. (A) Our green AF imaging studies showed growth stage-dependent changes of the basal green AF of the back’s skin and the ear’ skin of the mice. (B) The basal AF intensity of the back’s skin at the age of 1 d was significantly higher than that at the age of 1 w, 3 w and 10 w. N = 9; *; *p* < 0.05; ***; *p* < 0.001. (C) The basal AF intensity of the ear’s skin at the age of 10 w was significantly higher than that at the age of 3 w. N = 9; *; *p* < 0.05.

### 2. There are growth stage-dependent changes of the levels of K1 and K10 as well as skin’s green AF of the back and the ears of UVC-exposed C57BL/6 mice

We also determined K1 and K10 levels of UVC-exposed skin of the back and the ears of the mice at various age of the mice. Our Western blot assays showed that UVC induced significant decreases in the K1 levels of the back’s skin of the mice at the age of 1 d, 1 w, 3 w, and 10 w (Figs. 5A and 5B). UVC also induced significant decreases in the K10 levels of the back’s skin of the mice at the age of 3 w and 10 w (Figs. 5A and 5C). There were growth stage-dependent changes of the proteins of the back’s skin of the UVC-exposed mice (Figs. 5A, 5B and 5C). Our Western blot assays further showed that UVC induced significant decreases in the K1 levels of ear’s skin of the mice at the age of 3 w and 10 w (Figs. 5D and 5E). In contrast, UVC did not induce significant decreases in the K10 levels of the ear’s skin at the age of 3 w and 10 w (Figs. 5D and 5F). There were also growth stage-dependent changes of the proteins of the ear’s skin of the UVC-exposed mice (Figs. 5D, 5E and 5F).

**Fig.5.**
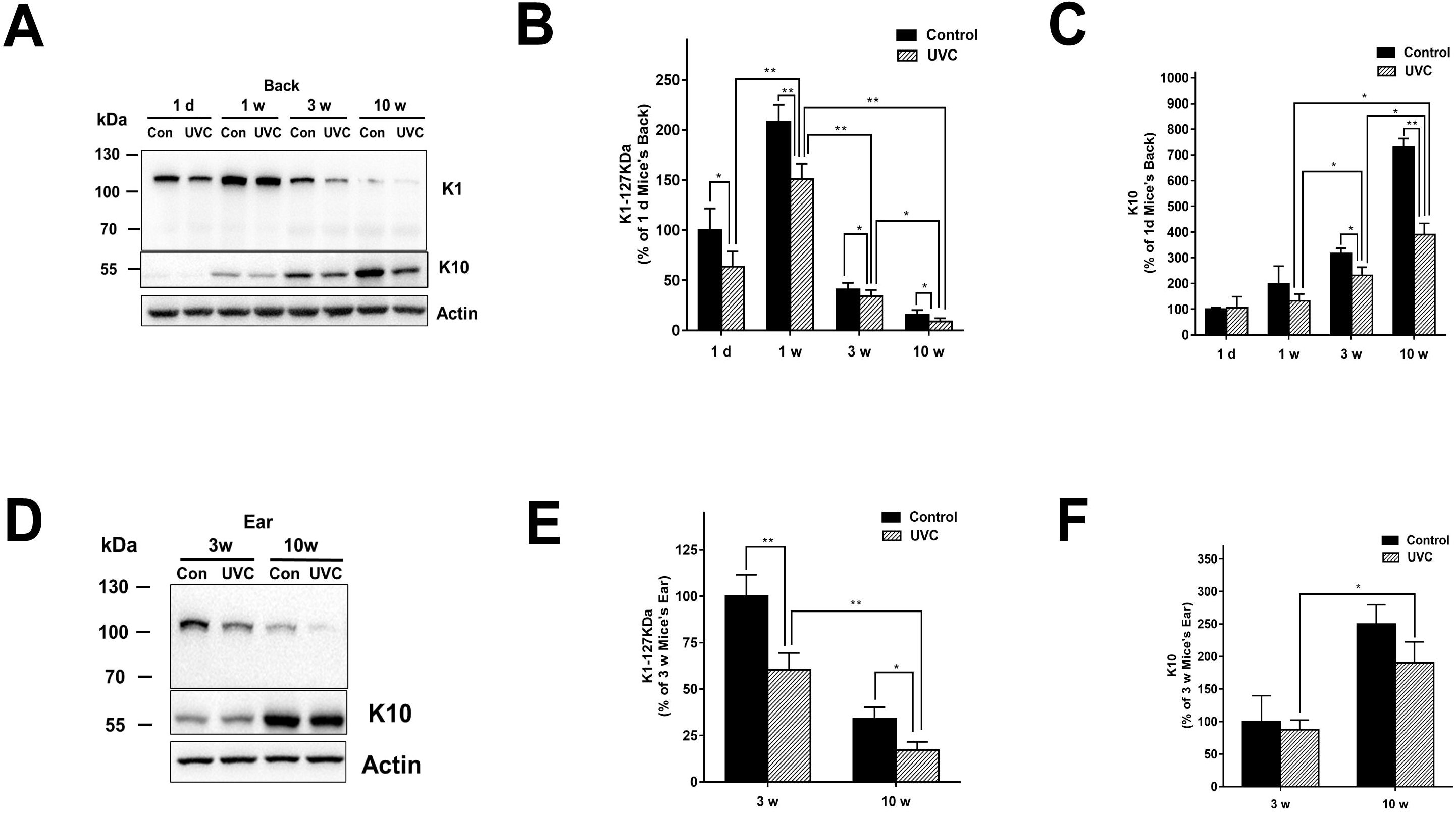
There were growth stage-dependent changes of the levels of K1 and K10 of the back’s skin and the ear’s skin of C57BL/6 mice after UVC exposures. (A) Western blot assays showed growth stage-dependent changes of the levels of K1 and K10 of the back’s skin of C57BL/6 mice at the age of 1 d, 1 w, 3 w, 10 w after UVC exposures. (B) Quantifications of the Western blot showed that UVC induced significant decreases in the K1 levels of the back’s skin of C57BL/6 mice at the age of 1 d, 1 w, 3 w, 10 w. N = 12; *; *p* < 0.05; **; *p* < 0.01. (C) Quantifications of the Western blot showed that UVC induced significant decreases in the K10 levels of the back’s skin of C57BL/6 mice at the age of 1 d, 1 w, 3 w, 10 w. N = 12; *; *p* < 0.05; **; *p* < 0.01. (D) Western blot assays showed growth stage-dependent changes of the levels of K1 and K10 of the ear’s skin of C57BL/6 mice at the age of 3 w and 10 w after UVC exposures. (E) Quantifications of the Western blot showed that UVC induced significant decreases in the K1 levels of the ear’s skin of C57BL/6 mice at the age of 3 w and 10 w. N = 12; *; *p* < 0.05; **; *p* < 0.01. (F) Quantifications of the Western blot showed that K10 levels of the ear’s skin of UVC-exposed C57BL/6 mice at the age of 3 w were significantly lower than those of the UVC-exposed mice at the age of 10 w. N = 12; *; *p* < 0.05.

Using anti-K1 antibody, our immunostaining assays on the back’s skin of the UVC-exposed mice at the age of 1d, 1 w, 3 w, and 10 w showed that the K1 levels changed in the similar pattern as those observed in our Western blot assays (Fig. 6A). Our immunostaining assays on the ear’s skin of the UVC-exposed mice at the age of 3 w and 10 w also showed that the K1 levels changed in the similar pattern as those observed in our Western blot assays (Fig. 6B).

**Figure 6.**
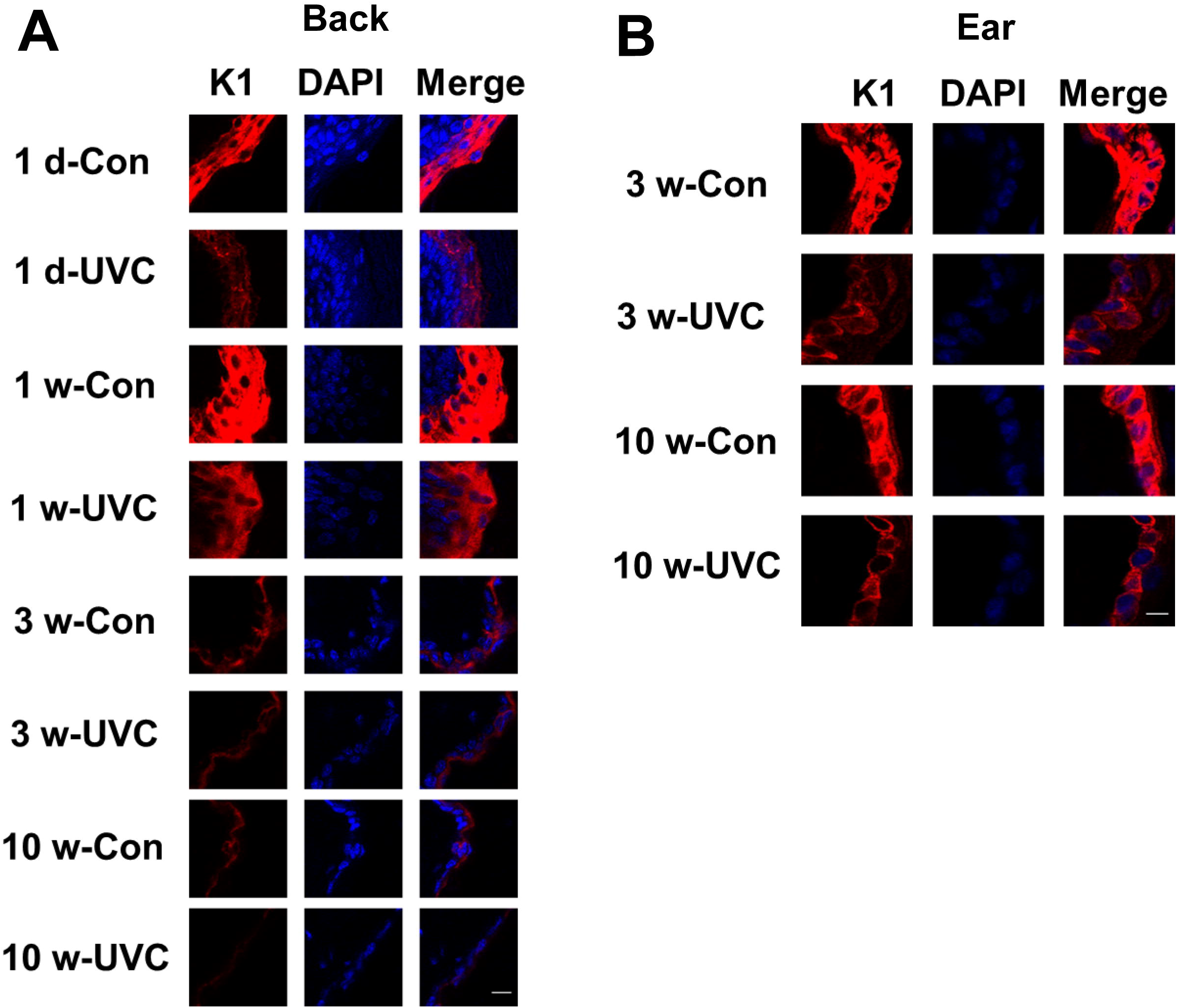
Immunostaining assays using anti-K1 antibody on the back’s skin and ear’s skin of the UVC-exposed mice at the various growth stages showed that both K1 and K10 levels changed in the similar patterns as those observed in our Western blot assays. (A) Our immunostaining assays on the back’s skin of the UVC-exposed mice at the age of 1d, 1 w, 3 w, and 10 w showed that the K1 levels changed in the similar patterns as those observed in our Western blot assays. N = 3 - 6. Scale bar = 5 μm. (B) Our immunostaining assays on the ear’s skin of the UVC-exposed mice at the age of 3 w and 10 w also showed that the K1 levels changed in the similar patterns as those observed in our Western blot assays. N = 3 - 6. Scale bar = 5 μm.

We also determined the green AF of the skin of mice’s back and ears at various ages after UVC exposures: UVC exposures induced significant increases in the green AF intensity of the back’s skin at the age of 1 d, 1 w, 3 w, and 10 w (Figs. 7A and 7B). UVC exposures also induced significant increases in the green AF intensity of the ear’s skin at the age of 3 w and 10 w (Figs. 7C and 7D). There were also growth stage-dependent changes of the UVC-induced green AF of both backs’ skin and the ear’s skin of the UVC-exposed mice (Figs. 7A, 7B, 7C and 7D).

**Figure 7.**
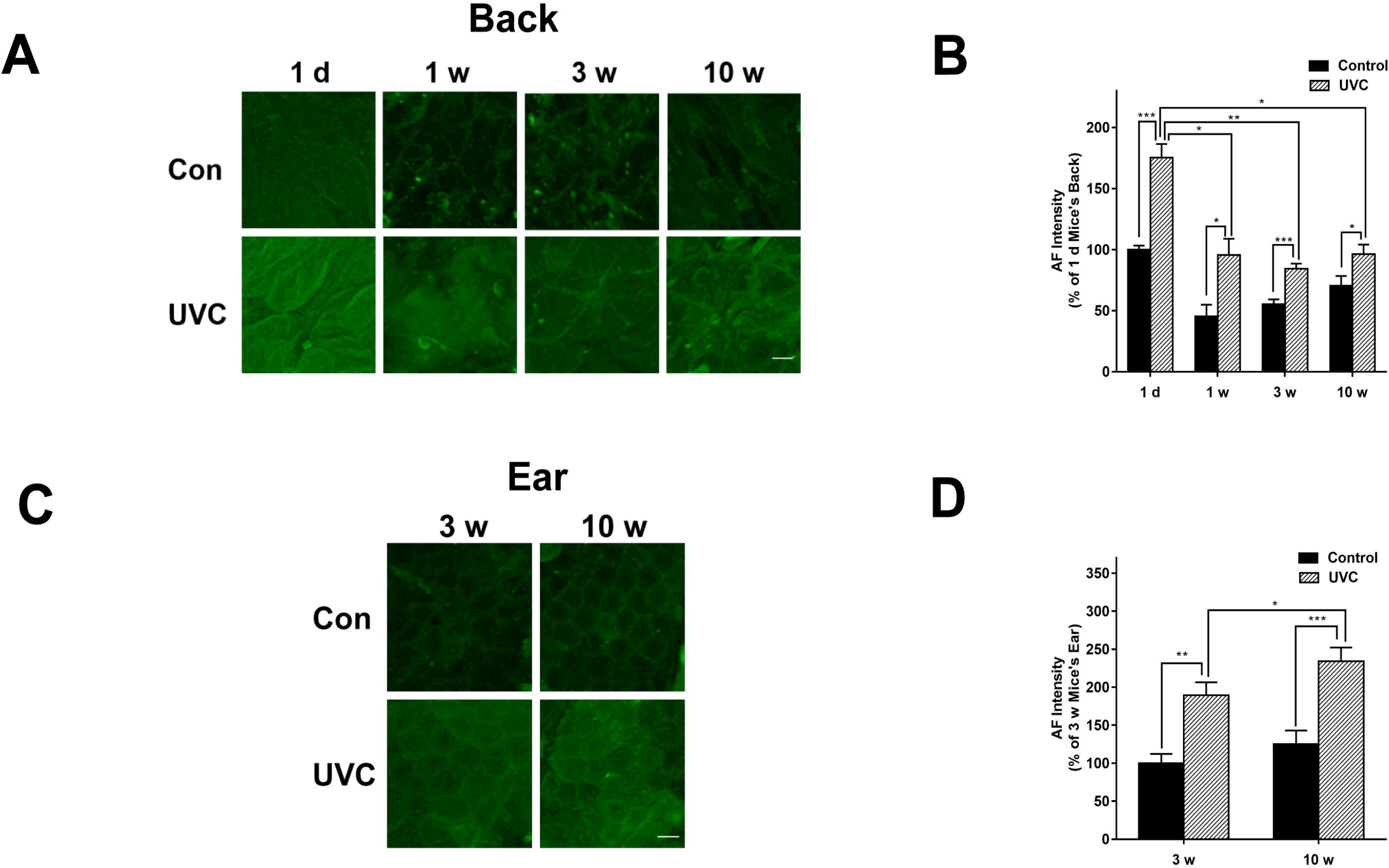
There were growth stage-dependent changes of the green AF of the back’s skin and the ear’ skin of the UVC-exposed mice. (A, B) UVC exposures induced significant increases in the green AF intensity of the back’s skin of the mice at the age of 1 d, 1 w, 3 w, and 10 w. Scale bar = 20 μm. N = 12. *; *p* < 0.05; **; *p* < 0.01;***; *p* < 0.001. (C, D) UVC exposures induced significant increases in the green AF intensity of the ear’s skin of the mice at the age of 3 w and 10 w. N = 12. *; *p* < 0.05;**; *p* < 0.01; ***; *p* < 0.001.

## Discussion

The major findings of our current study include: First, in the skin of both mice’s ears and back, there are growth stage-dependent changes of the levels of K1 and K10; second, in the skin of both ears and back of UVC-exposed mice, there are also growth stage-dependent changes of the levels of K1 and K10; third, there were growth stage-dependent changes of the green AF intensity of both back’s skin and the ear’s skin of the mice under basal conditions and after UVC exposures; and fourth, both K1 and K10 levels of the ear’s skin of the mice are remarkably different from those of the back’ skin.

Our study has indicated that K1 and K10 are major potential origins of the green AF induced by inflammation and oxidative stress ^[19]^. Our previous studies have indicated that the patients of each of the diseases we have studied, including AIS ^[20]^, myocardial infarction ^[21]^, stable coronary artery disease ^[21]^, Parkinson’s disease and lung cancer ^[23]^, had increased green AF in various locations of their skin, compared with age-matched healthy population. Our study has also indicated age-dependent changes of the skin’s green AF ^[24]^. However, there has been no information indicating if there are growth stage-dependent changes of K1 and K10.

In our current study, we used a mouse model to test our hypothesis that there are growth stage-dependent changes of the K1 and K10 levels, which has provided evidence supporting our hypothesis: In the skin of both mice’s ears and back, there are growth stage-dependent changes of K1 and K10 levels. These findings have suggested that the growth stage-dependent changes of the K1 and K10 levels may be an important mechanism underlying the age-dependent changes of the skin’s green AF of the healthy population. These findings have also indicated that we should use only age-matched healthy population as the healthy controls for the keratins’ AF-based diagnosis of diseases.

Our finding regarding the growth stage-dependent changes of the K1 and K10 levels is also of importance for understanding the pathological changes and mechanisms of multiple K1- and K10-associated diseases: For these diseases, certain pathological changes may be growth stage-dependent, which may result from the growth stage-dependent changes of the K1 and K10 levels in the skin. Future studies are needed to identify the growth stage-dependent pathological changes of these diseases.

Our study has also found that the levels of both K1 and K10 in the ear’ skin are remarkably different from those of the back’ skin. This finding has indicated that there may be location-dependent changes of K1 and K10 levels, which may be one of the potential factors accounting for the location-dependent changes of the skin’s AF for the multiple diseases. It is noteworthy that numerous biological studies on the skin used only one location of the skin for their studies ^[25, 26]^. Our finding has suggested that the findings from the study using only one location of the skin may not be applicable to the other locations of the skin. Since K1 and K10 can not only act as cytoskeletal proteins ^[27, 28]^, but also play significant roles in multiple biological processes such as inflammation ^[29–31]^, our study has suggested the location-dependence of the K1 and K10 levels may lead to significant location-dependence of multiple K1- and K10-associated biological properties.

Collectively, our study has provided first evidence showing growth stage-dependent and differential changes of the levels of K1 and K10 as well as skin’s green AF in the back and the ears of mice under basal conditions and after UVC irradiation. These findings are important for not only establishing the keratins’ AF-based method for non-invasive diagnosis of diseases, but also understanding the pathological mechanisms of K1 mutations- and K10 mutations-associated diseases.

## Acknowledgment

The authors would like to acknowledge the financial support by two research grants from a Major Special Program Grant of Shanghai Municipality (Grant # 2017SHZDZX01) (to W.Y.) and a Major Research Grant from the Scientific Committee of Shanghai Municipality #16JC1400502 (to W.Y.).

